# Spatiotemporal precise optical manipulation of intracellular molecular activities

**DOI:** 10.1101/2023.07.19.549752

**Authors:** Bin Dong, Shivam Mahapatra, Matthew G. Clark, Mark Carlsen, Karsten J. Mohn, Seohee Ma, Kent Brasseale, Grace Crim, Chi Zhang

## Abstract

Controlling chemical processes in live cells is a challenging task. The spatial heterogeneity of biochemical reactions in cells is often overlooked by conventional means of incubating cells with desired chemicals. A comprehensive understanding of spatially diverse biochemical processes requires precise control over molecular activities at the subcellular level. Herein, we develop a closed-loop optoelectronic control system that allows the manipulation of biomolecular activities in live cells at high spatiotemporal precision. Chemical-selective fluorescence signals are utilized to command lasers that trigger specific chemical reactions or control the activation of photoswitchable inhibitors at desired targets. We demonstrate the capability to selectively produce reactive oxygen species (ROS) solely at targeted organelles using blue light. Notably, the induction of ROS in the endoplasmic reticulum leads to a more pronounced disruption of tubulin polymerization and a reduction in green fluorescent protein signals, in comparison to that in lipid droplets. Moreover, when combined with a photoswitchable inhibitor, we selectively inhibit tubulin polymerization within subcellular compartments. This technology enables spatiotemporal control over chemical processes and drug activities, exclusively at desired targets, while minimizing undesired effects on non-targeted locations.

## Introduction

Biomolecular activities within cells exhibit intricate spatiotemporal heterogeneity. Advanced microscopy technologies have provided the means to visualize and study this heterogeneity in real-time and at high spatial resolution^1-5^. However, current methods of controlling chemical processes in live cells remain rudimentary and often fail to address this heterogeneity. The prevailing approach involves incubating cells with chemical compounds that interact with molecular targets. However, these interactions lack spatial precision, leading to potential off-target effects. The ability to precisely manipulate drug or protein activities in both space and time would provide valuable insights into biological functions and drug-target interactions. Laser-based optical control technologies offer the potential to achieve high spatiotemporal precision^6-9^. Laser beams can be focused to as small as hundreds of nanometers and delivered instantaneously to targets of interest. Existing optical control methods rely on microscopes to gather prior knowledge of the sample and visualize sample responses^10-12^. After image acquisition and analysis, laser interaction sites are manually selected using galvo mirrors or structured illumination methods^8,10-12^. These requirements do not allow real-time precise selection and optical manipulation of dynamic proteins or organelles in cells.

To address these challenges, we developed a real-time precision opto-control (RPOC) technology based on a closed-loop optoelectronic feedback system pair with a laser scanning microscope. This innovative system can detect chemical-specific optical signals in live cells, decide pixels of interaction in real-time, and manipulate chemical processes with high spatiotemporal precision solely at targets of interest. A gamut of continuous wave (CW) lasers enables simultaneous target selection, optical control, and monitoring of cellular responses. The incorporation of advanced comparator circuitry and acousto-optic modulators allows for active determination and precise commanding of separate laser wavelengths at desired pixels within the pixel dwell time. Importantly, optical control pixels are automatically selected based on the chemical signatures, eliminating the need for *a priori* knowledge of the sample. Using the RPOC technology, we successfully demonstrated selective induction of reactive oxygen species (ROS) in distinct organelles using a 405 nm laser. Quantitative analysis and laser dosage dependence results indicate that inducing ROS in the endoplasmic reticulum (ER), compared to lipid droplets (LDs), is associated with a more pronounced disruption of tubulin polymerization and a greater loss of green fluorescent protein signals. Furthermore, by coupling a photoswitchable inhibitor with RPOC, we achieve selective inhibition of tubulin polymerization in subcellular compartments. This capability enables the activation of inhibitors exclusively at desired targets while inactivating the same molecules at unwanted locations, providing precise spatiotemporal control over biomolecular activities. Our system offers a powerful tool for investigating cellular dynamics and unlocking new possibilities in drug development and cell biology.

## Results

### A precision opto-control technology for spatiotemporal manipulation of biomolecules

The real-time precision opto-control (RPOC) system is based on a laser scanning confocal fluorescence microscope that enables highly specific chemical detection and imaging. As illustrated in **Fig. 1a**, four CW lasers facilitate signal excitation and optical control. The system encompasses three key components: chemical target selection, optical manipulation, and readout of cell responses. Both the 473nm laser and 589nm laser are utilized to excite fluorescent labels for the selection of chemical targets for optical manipulation. The 473 nm laser, which matches the excitation wavelength of green fluorescent proteins (GFPs), functions as the readout laser for cell responses. The 532 and 405 nm lasers are optical switches of chemical processes at desired locations. Two photomultiplier tube (PMT) detectors are set up in confocal configurations. One PMT is utilized to detect targets for optical manipulation, while the other is designated as a readout detector. Fluorescent signals from the chemical selection PMT are sent to a comparator circuit for decision-making. This circuit is responsible for analyzing the fluorescent signals and triggering acousto-optic modulators (AOMs) in the opto-control laser paths accordingly. If the intensity of the fluorescent signal exceeds a respective threshold, a ‘1’ Transistor-Transistor Logic (TTL) command is sent to the AOMs to deflect the control laser to the 1^st^-order direction to spatially combine with other laser beams. Otherwise, a ‘0’ TTL command on the AOM prevents the delivery of the control laser to the sample. The two AOMs can be controlled separately based on the chemical signatures of the sample at each pixel. A detailed illustration of the comparator circuit is discussed in the Supplementary information **(Supplementary Note 1, Supplementary Fig. 1)**. The flow of the CW-RPOC system is shown in **Fig. 1b**. The chemical target selection, decision-making, and opto-control are performed within a single pixel, paired with simultaneous real-time imaging of cells from the readout channel. An active pixel (APX) is defined as the pixel of which the opto-control laser is activated ^13^. APX can be visualized by splitting the output from the comparator circuit or by using photodiodes collecting small fractions of the opto-control lasers.

**Figure 1.**
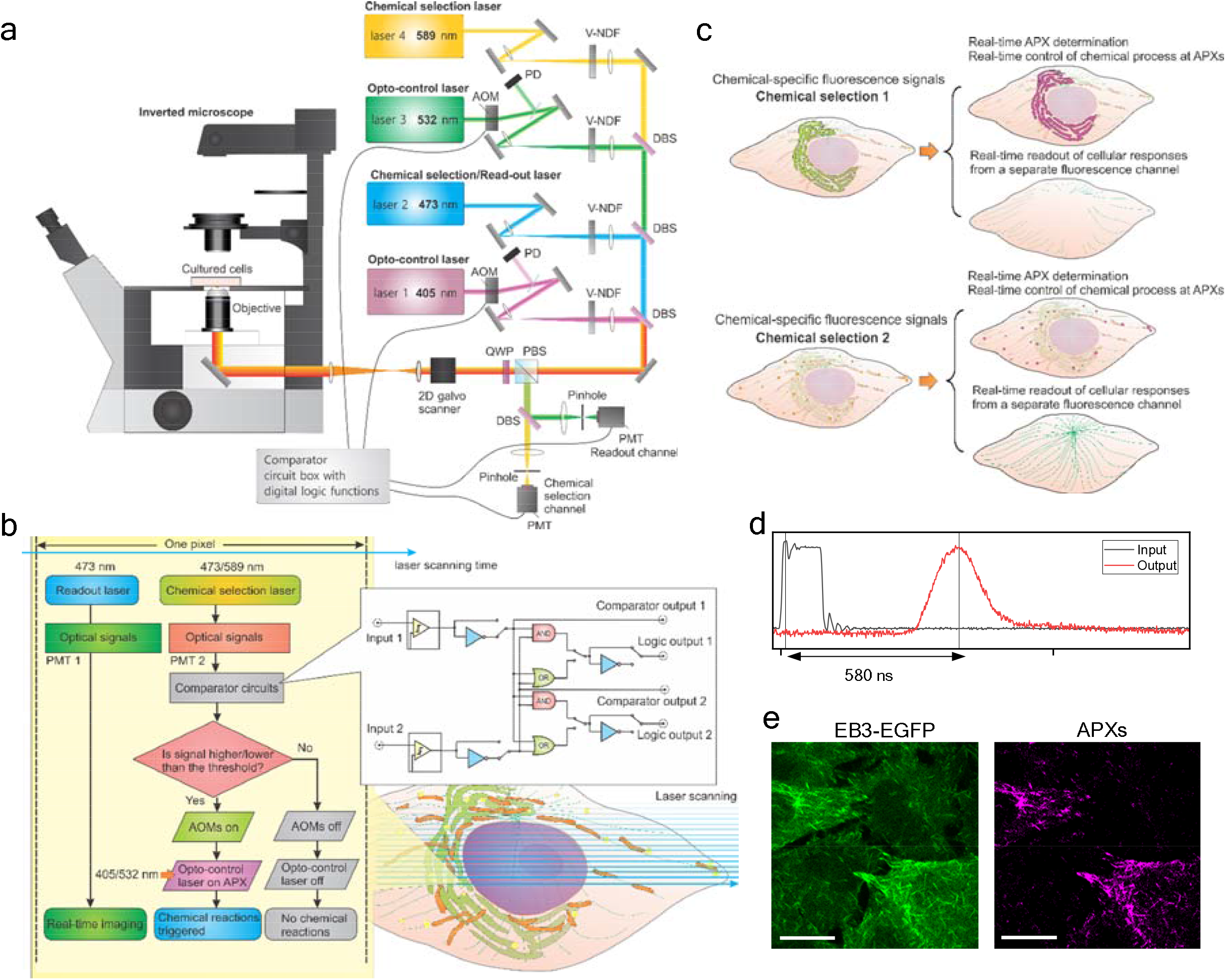
The CW-RPOC platform for precision opto-control. (a) The optical configuration of the CW-RPOC system. AOM, acousto-optic modulator; DBS, dichroic beamsplitter; PBS, polarization beamsplitter; V-NDF, variable neutral density filter; QWP, quarter-wave plate; PMT, photomultiplier tube. (b) The workflow of CW-RPOC at a single pixel and the comparator circuitry. (c) An illustration of selective chemical detection, active pixel (APX) determination, and real-time monitoring of cell responses using the CW-RPOC system. (d) The response time of the RPOC system for the 405 nm laser. (e) A fluorescence image of EB3-EGFP signals from HeLa cells and the corresponding APXs selected using the fluorescence signals.

APX can be specifically chosen for desired molecular targets in cells. For example, as shown in **Fig. 1c**, fluorescent labels can be used to exclusively select APX either on ER or LDs. The readout signal from GFP allows simultaneous monitoring of cellular responses to the optical control targeting selected organelles. The opto-control laser power, pixel dwell time, and wavelength can be separately tuned to explore the impact of laser dosage and wavelength on the manipulation of cellular activities. The response time of the feedback loop, as measured to be approximately 580 ns for the 405 nm laser (**Fig. 1d, Supplementary Note 2, and Supplementary Fig. 4**), is significantly shorter than the 10-20 μs pixel dwell time. This rapid response time ensures tracking and active control of dynamic molecular targets within cells. **Fig. 1e** displays fluorescence signals from enhanced GFP (EGFP) conjugated with end-binding protein 3 (EB3) in HeLa cells, as well as APXs selected by the EGFP signals. EB3 binds to the plus end of the microtubule during tubulin polymerization^14^. The EB3-EGFP signals reveal highly dynamic tubulin polymerization processes in HeLa cells and highlight RPOC for tracking moving targets for opto-control (**Supplementary Videos 1.1-1.3**). The system exhibits a diffraction-limited spatial resolution of approximately 300 nm, with a similar spatial precision for opto-control (**Supplementary Note 3, Supplementary Fig. 5**). The pixel size, in the oversampling condition, is typically around 100 nm.

### Inducing reactive oxygen species precisely by blue light at selected organelles

In the first application, we explore using CW-RPOC to precisely induce ROS by blue light only at desired organelles and study cell responses to such organelle-associated ROS generation. ROS are highly reactive molecules containing oxygen atoms with an unpaired electron. While ROS are natural byproducts of metabolic processes and are well-regulated in live cells^15-17^, increased ROS generation induced by pathological transitions or external stimuli can cause oxidative stress that significantly affects cell functions^18-20^. Depending upon the source of ROS generation, different organelles have diverse impacts on cellular pathways. For instance, Zeng and coworkers used organelle-targeting photosensitizers and demonstrated that pyroptosis is stimulated most efficiently by ER-targeted photodynamic therapy^21^. This makes it crucial to understand organelle-specific responses to ROS generation, especially when ER is stimulated. It is known that short-wavelength blue light can induce ROS in cells^22,23^. However, existing methods of ROS generation in cells lack spatiotemporal control, making it hard to understand the impact of ROS associated with different subcellular targets such as ER or LDs. With the CW-RPOC system, we achieve subcellular-precise ROS generation solely on the targets of interest and quantify cell response to dosage-dependent ROS associated with different organelles.

HeLa cells with a stable EB3-EGFP expression are used for the study. Fluorescent organelle markers are excited by either the 589 or 473 nm laser to locate organelles of interest as opto-control targets. The 405 nm laser is used to directly induce ROS at targets by CW-RPOC. The EGFP signals, excited by the 473 nm laser, are used to visualize tubulin polymerization dynamics and quantify cell responses to localized ROS generation. Using ER-Tracker, APXs can be selected only at ER locations. The ER-Tracker signals, APXs, and EB3-EBFP fluorescence images during RPOC are shown in **Fig. 2a**. A significant EB3-EGFP signal decrease is observed when a 240 μW 405 nm laser is shone on ER with a pixel dwell time of 20 μs (**Fig. 2a**). Such a decrease is not observed in the control group (no APX) and treating ER with 150 μW 405 nm laser (**Fig. 2a**). When selecting APX on LDs using BODIPY LD labeling and applying 240 μW 405 nm laser uniquely at LDs, no considerable EB3-EGFP signal decrease is detected (**Fig. 2a**). Averaged time-dependent EB3-EGFP fluorescence signals from cells in each condition are normalized to compare trends in intensity change (**Fig. 2b**). Such curves are fit using exponential functions to obtain the decay time constants of each condition (**Fig. 2c, d**). The results reveal that a strong EB3-EGFP signal decrease is associated with 240 μW 405 nm interacting with ER, not LDs. Aside from the EGFP signal decrease, the 240 μW 405 nm laser interaction with ER causes a faster disruption of cellular tubulin polymerization compared to other cases, which can be seen in **Supplementary Video 2**. It is well-documented that blue light can induce ROS that impact cellular functions^22,23^. Our additional studies by treating cells using different laser wavelengths confirm that 405 nm laser disrupts tubulin polymerization in live cells (**Supplementary Note 4, Supplementary Fig. 6**). It was reported that ROS can decrease microtubule growth^24^. ROS on ER can potentially cause calcium leak that hampers microtubule polymerization and cellular GFP signals^8,25^. Our results indicate that cell response to localized blue light treatment is organelle dependent. Compared to LDs, ER is a more responsible target for blue light induced EGFP signal loss and tubulin dynamics disruption. Light-induced ROS on ER is more detrimental to cells than on LDs.

**Figure 2.**
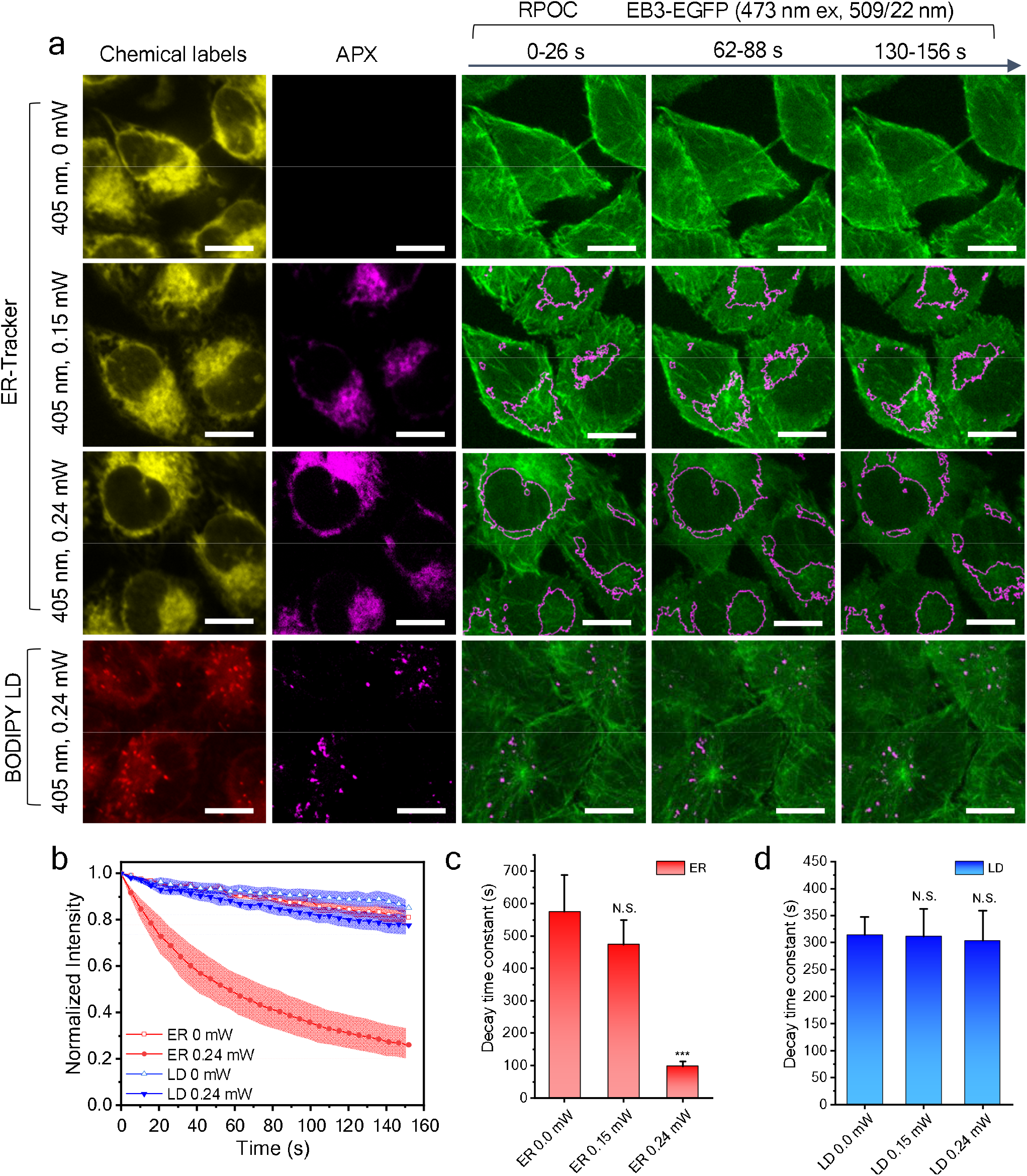
Organelle-dependent ROS generation by 405 nm laser using CW-RPOC. (a) Representation of chemical selection channels, active pixels (APX) selected, and time-dependent EB3-EGFP signals from the readout channel. In the first three rows, ER-are labeled using ER-Tracker. In the fourth row, lipid droplets (LDs) are labeled using BODIPY. To highlight ER-selected APX areas, outlines of the APXs are overlayed with EGFP signals. (b) Time-dependent EB3-EGFP signal change for cells in different treatment conditions. (c, d) The decay time constants of EB3-EGFP signals for cells when APXs are selected on ER (c) and LDs (d). Scale bars are 10 μm. For statistical analysis, *** p<0.001, N.S. p>0.05.

**Fig. 2a** also implies that cell responses to localized ROS generation are laser-dosage-dependent. To correlate EGFP signal changes to laser dosage, a quantitative analysis method is developed. The laser dosage can be quantified using APX intensity. As shown in **Fig. 3a**, APX intensity measures the 405 nm laser power multiplying the total time of the fluorescence signals exceeding the intensity threshold in the APX. If the target fluorescence signal is strong, the signal would exceed the threshold for a longer time, resulting in higher APX intensity. The maximum APX intensity is measured when the fluorescence signal is above the threshold throughout the entire APX dwell time. **Fig. 3b** shows example APX images and the corresponding APX intensity profiles at dash lines when ER and LDs are selected. Integration of APX intensity in a cell quantifies the dosage of 405 nm laser for ROS generation. The quantification procedures are detailed in **Materials and Methods**. ER boundaries of two HeLa cells are shown in **Fig. 3c**. The laser dosage of cell 1 is around 123 μJ, which is much lower than 296 μJ for cell 2. Despite both being treated on ER, cell 1 and cell 2 show different intensity decay profiles due to the laser dosage difference (**Fig. 3d**). A higher laser dosage tends to induce a faster decay of EB3-EGFP signals (**Fig. 3d**). We quantify 405 nm laser dosage for ER and LD in each cell and plot their correlations with the corresponding EGFP signal decay (**Fig. 3e**). The difference in such correlations for ER and LD indicates 405 nm laser-induced ROS have different impacts on different organelles. Note that for the untreated case (0 dosage), LD-labeled cells have a faster EGFP signal decay compared to ER-labeled cells. This is likely due to the higher cytotoxicity and functional perturbation induced by BODIPY compared to ER-Tracker. However, as the 405 nm laser dosage increases, the same laser dosage on ER causes a faster EGFP signal decrease than on LDs. Such an escalated EGFP signal loss is likely mediated by an additional pathway associated with ER, as illustrated in **Fig. 3f**. It has been reported that an excess amount of ROS in ER can cause the leaking of calcium into the cytosol. Such released calcium can induce mitochondrial dysfunction and a cascade of apoptotic events ^8,26^. This ER-associated ROS likely induces the acceleration of EGFP signal loss. This study highlights the capability of inducing ROS with blue light at selected organelles and simultaneously monitoring cellular responses to localized ROS generation.

**Figure 3.**
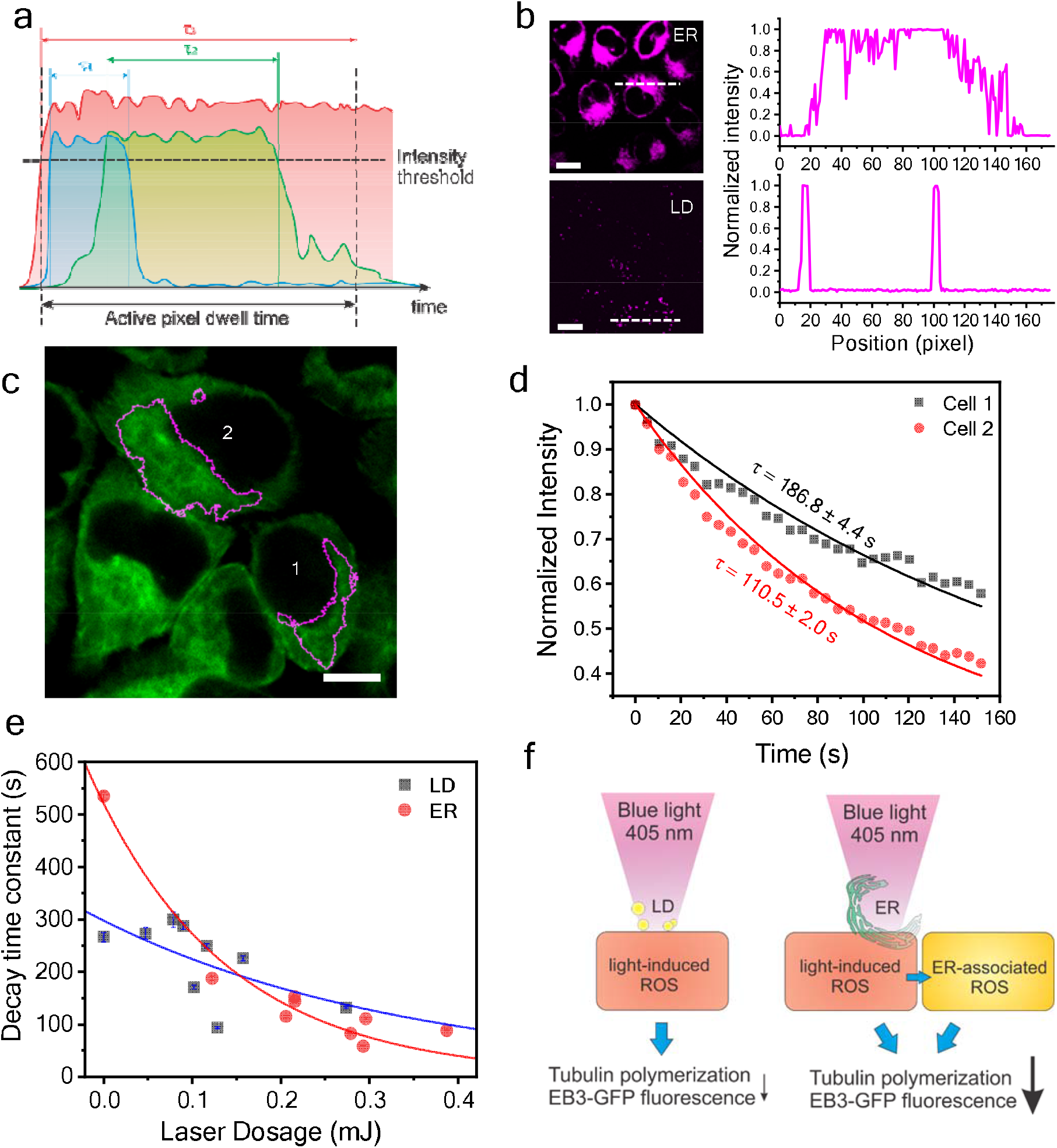
Laser-dosage-dependent cell responses to organelle-specific ROS. (a) An illustration of different optical signals in an APX and the corresponding time (τ_1_, τ_2_, τ_3_) when signal levels are above the intensity threshold. (b) Example images of APX on ER and LD and the corresponding APX line profiles. (c) EB3-EGFP signals of HeLa cells and the outlines of ER-APXs selected for two cells. (d) Time-dependent EB3-EGFP signal changes for cells 1 and 2 in panel c. Dots are experimental results; lines are fitting curves using an exponential decay function. (e) EB3-EGFP signal decay time constant plotted against the 405 nm laser dosage for LD and ER. Dots are experimental results; lines are fitting curves using an exponential decay function. (f) A schematic representation of different effects of 405 nm laser interacting with ER and LD in live cells. Blue light treatment on ER likely induces additional ER-associated ROS causing a greater decrease of EB3-EGFP signals and tubulin polymerization. Scale bars are 10 μm.

### Precise inhibition of biomolecular activities at subcellular targets

Apart from directly inducing chemical changes by lasers, RPOC can selectively control the activities of photoswitchable compounds to inhibit chemical processes exclusively at subcellular targets. A photoswitchable tubulin polymerization inhibitor photostatin (PST-1) was synthesized as previously reported^13,27^. The photoswitchable active *cis-* and inactive *trans-*isomers of PST-1 are shown in **Fig. 4a**. Selective photoactivation of PST-1 allows for inhibition of tubulin polymerization at selected targets in cells (**Fig. 4a**). To rule out the potential impact of blue-light induced ROS, 30 μW of 405 nm laser is used. Such a laser dosage does not induce detectable decreases in EB3-EGFP signals. Furthermore, the inhibition of tubulin polymerization does not decrease EB3-EGFP signals integrated over a whole cell (**Supplementary Note 4, Supplementary Fig. 6**). This is because the depolymerized EB3 molecules are released into cytosols and remain fluorescent.

**Figure 4.**
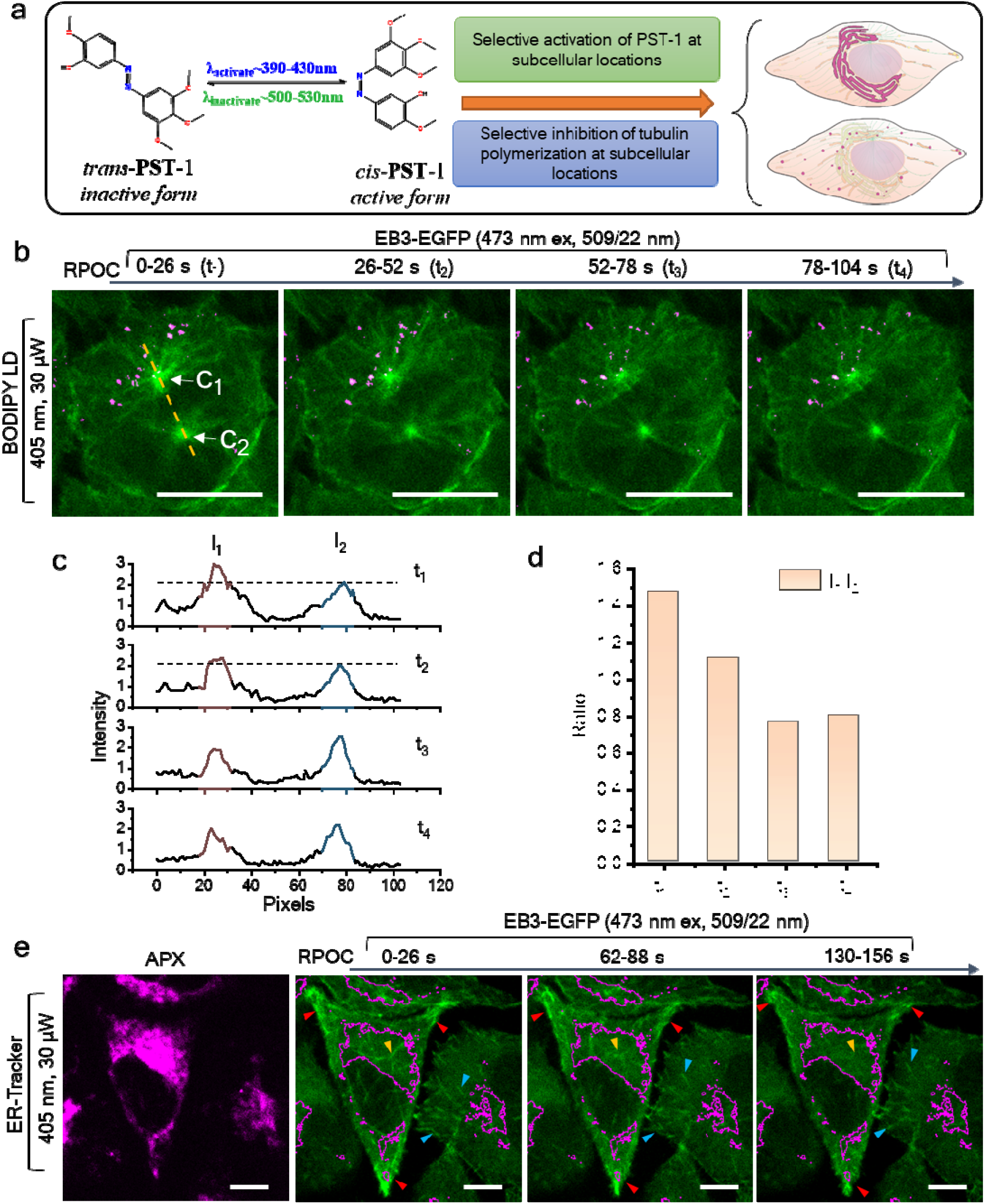
Precision inhibition of tubulin polymerization at subcellular compartments. (a) Interconversion of *trans*-PST-1 and *cis*-PST-1 upon stimulation by blue and green laser wavelengths. An illustration of selective activation of PST-1 only on ER and around LDs by RPOC. (b) EB3-EGFP signals over time with APX displayed in magenta. Cells are treated with inactive PST-1 which is only activated at APXs. APXs are associated with centrosome C_1_ instead of C_2_. (c) The change in the intensity of EB3-EGFP signals along the dashed line in panel b for two centrosomes. (d) The ratio of *I*_*1*_ over *I*_*2*_ as a function of RPOC activation time. (e) APX selected by ER-Tracker and EB3-EGFP signals at different time points when PST-1 is selectively activated on ER. The ER APX areas are outlined in EB3-EGFP images. Scale bars are 10 μm.

Due to the highly dynamic signature of EB3-EGFP signals, it is hard to quantify the inhibition of tubulin at single LDs. Instead, we select centrosomes, which are EB3 abundant and are constantly bright in fluorescent images, to quantify the inhibition process. **Fig. 4b** represents a field of view of HeLa cells with two centrosomes detected. Cells are incubated with PST-1 in the inactive form before RPOC. LDs are labeled using BODIPY and PST-1 molecules are activated at the LD areas, which are close to and present on one of the centrosomes (C_1_). The other centrosome (C_2_), on the other hand, is distant from LD APXs. Time-lapse EB3-EGFP intensity shows a decrease in C_1_ together with no change for C_2_ during RPOC (**Fig. 4b, c, Supplementary Video 3**). The ratio of centrosome intensities I_1_/I_2_ further highlights such a change, which is due to selective inhibition of tubulin polymerization at C_1_ but not C_2_. These results demonstrate that even inside a single cell, RPOC can inhibit biomolecular activities at subcellular compartments and visualize cell responses simultaneously.

Using ER-Tracker to select APX on ER, RPOC can activate PST-1 and inhibit tubulin polymerization only in the ER area, as shown in **Fig. 4e**. Compare with the cell protrusion area (red arrows), tubulin dynamics on ER (yellow arrows) are more notably decrease (**Supplementary Video 4**). Note that when only using blue light for PST-1 activation, the activated compound can gradually diffuse into areas outside APX, and eventually lead to inhibition of tubulin dynamics of the whole cell, as illustrated in **Fig. 5a**. To prevent such active drug diffusion, 532 nm laser can be utilized to inactivate PST-1 outside APXs of 405 nm laser (**Fig. 5b**). This function can be achieved by the CW-RPOC system since the 405 nm and 532 nm lasers are separately controlled by two AOMs and different channels of the comparator circuitry. Here, 1.2 μW 405 nm and 1.6 μW 532 nm lasers are used for optical manipulation. **Fig. 5c** shows EB3 signals, APXs from two lasers, and time-lapse EB3 signal changes over time. In the top panels, PST-1 is mostly activated in the EB3-dense area (mostly centrosome) and inactivated elsewhere, while the bottom panels reverse the activation/inactivation locations. Comparing EB3-EGFP signals from centrosomes, we found that PST-1 activation on centrosomes by blue light shows a more pronounced signal decrease compared to that is inactivated by green light (**Fig. 5d, Supplementary Videos 5, 6**). Such a change is also observed in the cytosol areas with low EB3-EGFP signals (**Fig. 5e**). The decrease of EGFP signals during 532 nm laser treatment in cells is induced by the photobleaching of EGFP by the 473 nm laser. The inactivation of PST-1 by 532 nm laser increases the concentration of EB3 in the less polymerized locations. This process compensates for the photobleaching of EGFP, resulting in a flat plateau in the first 40 s in **Fig. 5e**. For EB3-dense centrosomes, such compensation is more efficient due to the lower dosage and interaction area of the 532 nm laser, giving a much slower signal decay as shown in **Fig. 5d**. Due to the selection of a much smaller area, the centrosome EGFP signal changes are more susceptible to the impact of tubulin dynamics, resulting in higher fluctuations in the intensity change (**Fig. 5d**). This study exemplifies that RPOC can selectively activate photoswitchable inhibitors and enable subcellular site-specific inhibition of biomolecular processes at high spatiotemporal precision.

**Figure 5.**
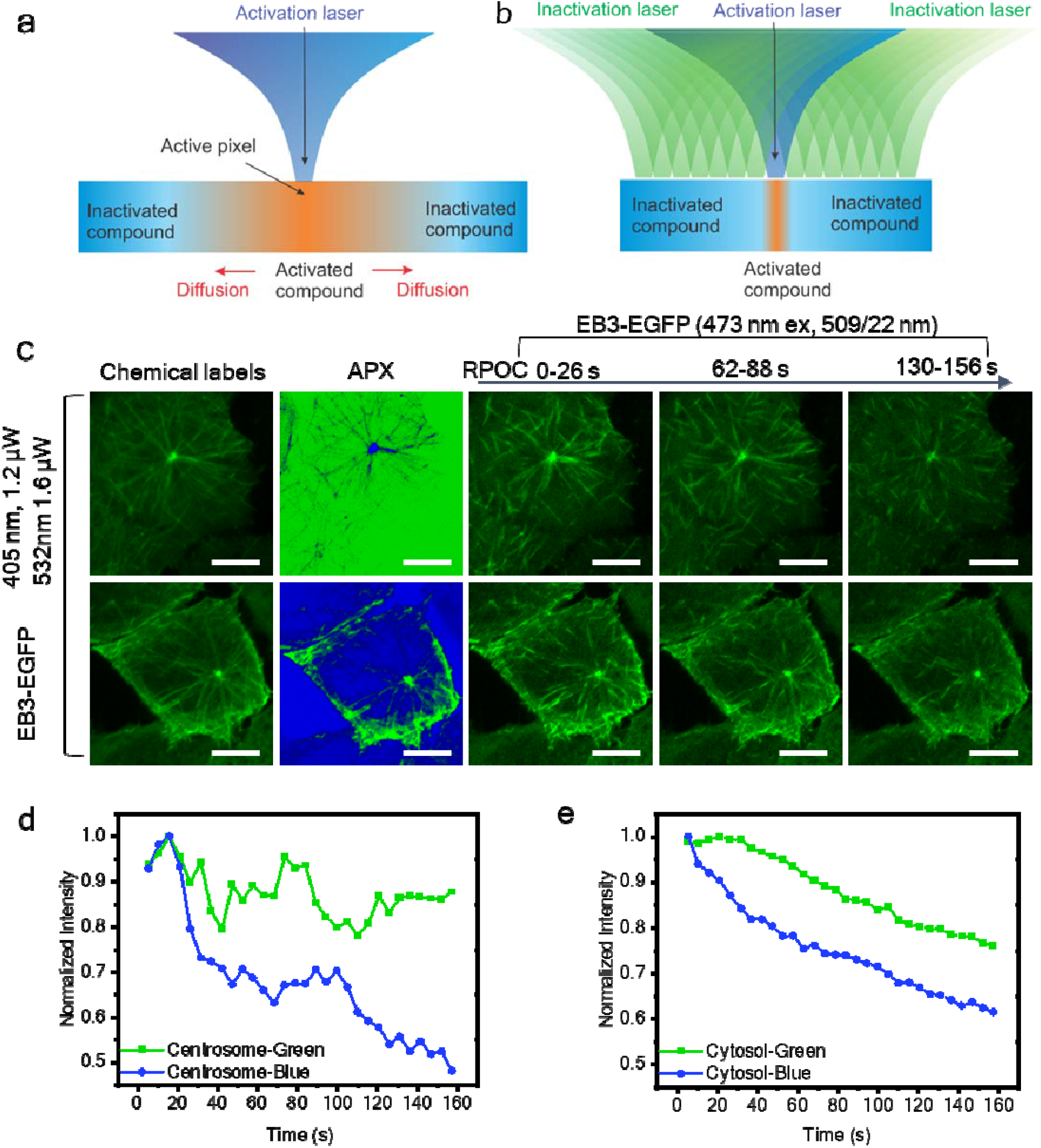
Subcellular control of photoswitchable inhibitor activities. (a) A schematic representation of diffusion of an activated photoswitchable compound to other subcellular locations in the absence of an inactivation laser. (b) A schematic representation of the inactivation of an activated compound in all locations other than desired target sites using the inactivation laser. (c) Chemical labels, APXs, and EB3-EGFP signals during selective activation/inactivation of PST-1 at different subcellular locations. Top row: inactivation of PST-1 only at the centrosome while activating it elsewhere. Bottom row: activation of PST-1 only at the centrosome and the cell boundary while inactivating it elsewhere. Green in the APX channel indicates APX for 532 nm, while blue in the APX channel indicates APX for 405 nm. (d) EB3-EGFP signals at centrosomes when the centrosome is treated with blue or green light. Cells are incubated with inactivated PST-1 before RPOC. (e) Similar to panel d but for cytosol areas excluding centrosomes and other EB3-dense areas. Scale bars are 10 μm.

## Discussion

Conventionally, it is impossible to precisely control chemical reactions and enzyme activities in cells at submicron precision. RPOC provides a new approach using lasers to manipulate biomolecular processes at high spatial precision and in real-time. We demonstrate precision induction of ROS at targeted organelles and correlate laser dosages to cellular response instantaneously. Furthermore, subcellular-localized inhibition of tubulin polymerization at selected organelles is achieved. RPOC does not require *a priori* knowledge of chemical targets and accomplishes target detection, decision-making, and opto-control simultaneously. This is especially advantageous to track and manipulate targets with highly dynamic signatures such as LDs, mitochondria, lysosomes, tubulin, actin, and other proteins with high mobility.

Compared to femtosecond lasers for signal excitation^13^, CW-RPOC is based on CW lasers that are cost-effective and compact. In this work, only 405 nm and 532 nm lasers are used for optical manipulation. Applying more lasers and adding more AOMs would allow more fluorescent labels for chemical detection and additional laser wavelengths for opto-control. Aside from EGFP, other fluorescent proteins transfected to cells or fluorescent labels can be applied as the readout for RPOC. The comparator circuitry is made with common electronics and can be easily fabricated. The laser scanning and CW-laser configuration make CW-RPOC intrinsically compatible with laser-scanning confocal fluorescent microscopes for broad applications in biological sciences.

We demonstrated precision control of tubulin dynamics in live cells using PST-1. Photoswitchable inhibitors of other cytoskeletal proteins or cellular enzymes can be paired with CW-RPOC for precision manipulation of a wide arrange of biomolecules in cells^28-34^. This approach would lead to new insights into cellular function through location-specific manipulation of biomolecular activities, subcellular activation of drugs, and precision-controlled release.

By shedding light on previously inaccessible aspects of cellular processes, RPOC opens up new avenues for understanding drug-target interactions, exploring site-specific biomolecular activities, and facilitating precise control over chemical release. On a larger scale, RPOC might bring about photodynamic therapy and treatment strategies with higher spatial and chemical precision and lower side effects.

## Materials and Methods

### The CW-RPOC technology

A detailed schematic of the RPOC system is shown in **Fig. 1**. The system incorporates four diode-pumped solid-state lasers (CNI Laser), which are collinearly combined using three dichroic beamsplitters (DMLP567, DMLP505, DMLP425, Thorlabs). Each laser is collimated by a pair of lenses, and the laser power is adjusted using a variable neutral density filter (V-NDF, 54-081, Edmund Optics). Laser 1 (λ_exc_ = 405nm, 100mW) and Laser 3 (λ_exc_ = 532nm, 100mW) are dedicated to optical control and are independently controlled by two acousto-optic modulators (AOMs, M1205-P80L-0.5 with 532B-2 driver, Isomet). A small portion of both Laser 1 and Laser 3 is directed to separate photodiode detectors (PDA10A2, Thorlabs) for visualization of APX. Laser 2 (λ_exc_ = 473nm, 100mW) is used for exciting fluorescent proteins in living cells. Laser 4 (λ_exc_ = 589nm, 100mW) is employed for exciting chemical targets for opto-control. The laser beams are expanded separately by different lens pairs to approximately 2.5 mm in diameter. For Lasers 1 and 3, the lenses are placed after AOMs. The intensities of all lasers can be separately adjusted by variable neutral density filters (53-212, Edmund Optics). The combined beams pass through a polarizing beamsplitter (PBS251, Thorlabs) and a quarter-wave plate before entering a 2D galvo scanner set (Saturn-5 system, ScannerMAX). The laser beams are further expanded by a pair of lenses to fill the entrance of a water-immersion objective lens (UPlanSApo-S, 60X, NA = 1.20, Olympus) mounted on an inverted microscope (Olympus IX73). Sample positioning is achieved using a 3D translational stage (H117 with Motor Focus Drive and ProScan III system, Prior).

The back-reflected fluorescence signals are directed by the polarizing beamsplitter to two photomultiplier tubes (PMT, H7422-40, Hamamatsu). Confocal detections are enabled by placing two pinholes (P300HK, Thorlabs) at the sample conjugate focal plane. The fluorescence signals from EGFP are detected by the PMT in the readout channel, which is equipped with a bandpass filter (FF01-509/22, Semrock). The chemical targets are detected by the PMT in the chemical selection channel, which utilizes a bandpass filter (ET642/80m, Chroma Technology Corporation). These two channels are separated by a long-pass dichroic beam splitter (FF552-Di02, Semrock). The PMT output currents from both channels are converted to voltage and amplified using two preamplifiers (PMT4V3, Advanced Research Instruments Corporation) before being sent to the data acquisition system (PCIe-6363 paired with BNC-2110, National Instruments). A custom multichannel image acquisition software based on LabVIEW is employed for image display, saving, and APX selection. A two-channel comparator circuit box is designed to achieve APX selection and closed-loop feedback control. The detailed design and functions of the comparator circuitry can be found in **Supplementary Note 1**. The outputs in TTL format control the two AOMs separately. This box allows for the selection of APX from two different lasers (405 nm and 532 nm) based on the fluorescence channels either separately or in a logically combined manner.

### Laser dosage measurement

The APX signals from the 405 nm laser, which are directly captured by a photodiode, provide real-time feedback on the laser dosage for ROS generation. If the fluorescence signal is above the threshold for the entire pixel dwell time, the maximum APX intensity *I*_*max*_ is measured, which can be achieved at APX saturations during RPOC or constantly turning on the AOM using the comparator circuit box. *I*_*max*_ is used to normalize the laser dosage on the sample.

For the time-lapse analysis performed in this study, multiple frames are acquired during RPOC. To quantify the laser dosage for the entire interaction time for a cell, the EB3-EGFP signals in the readout channel are used to manually outline individual cells. The cell mask is then projected to the averaged APX image (over acquired *N* frames) for quantification. The average intensity *I*_*ave*_ and mask area *A* are measured for each cell. An area outside of the cell in the average APX image is selected to measure the background intensity *I*_*bg*_. The opto-control laser at the sample has an average power of *P*. The pixel dwell time for RPOC and imaging is *T*. The laser dosage for each cell is calculated to be

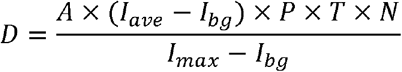

with a unit of J. All image processing, mask selection, and quantification are performed using ImageJ built-in functions.

### Quantitative image analysis

Pseudo-color labeled images of chemical labels, APXs, and EB3-EGFP are processed using ImageJ. In **Fig. 2**, EB3-EGFP images are averaged from 0-26 s, 62-88 s, and 130-156 s for display. Chemical labels and APX images are averaged from 0-156 s for display. To obtain time-dependent EB3-EGFP signal changes, single cells are manually outlined, and fluorescence signals gated in each cell are integrated for each frame. The integrated fluorescence intensity for each cell is plotted as a function of time for optical manipulation. The fluorescence intensity decay curves are subtracted by background intensities and normalized by dividing the maximum value. The normalized intensity decay curves can be fit using an exponential decay function:

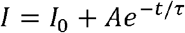

We used the decay curve with the fastest time constant to obtain *I*_*0*_ and *A*, which are maintained constant for the fitting of other conditions. The decay time constant r is compared for different conditions in **Fig. 2, 3, 5**. For the student’s t-test in **Fig. 2**, five cells are quantified in each condition.

APX intensity profiles in **Fig. 3 and 4** are plotted using an ImageJ built-in function.

### Cell preparation

HeLa Kyoto EB3-EGFP cells are purchased from Biohippo. Cells are cultured in Dulbecco’s Modified Eagle Medium (DMEM, ATCC) with 10% fetal bovine serum (FBS, ATCC) and 1% penicillin/streptomycin (Thermofisher Scientific). The cells are seeded in glass-bottom dishes (MatTek Life Sciences) with 2□mL culture medium and then incubated in a CO_2_ incubator at 37□°C and 5% CO_2_ concentration. Cells are grown to about 50% confluency and are directly used for live-cell imaging.

### PST-1 and cell treatment with PST-1

The synthesis of PST-1 can be found in a previous publication^13^. PST-1 is dissolved in dimethylsulfoxide at 2 mM concentration to prepare the stock solution. Before treating the cells, the PST-1 stock solution is treated with a 532 nm laser (CNI laser) for 5 s to convert PST-1 into the *trans* inactivated form. Cells are treated with inactivated PST-1 at a final concentration of 4□μM for 15□min before RPOC. The 405 nm laser (CNI laser) in the CW-RPOC system is used to selectively activate PST-1 solely at desired locations by CW-RPOC.

### Fluorescent labeling of ER and LDs

HeLa Kyoto EB3-EGFP cells are first seeded in glass-bottom dishes and cultured overnight to reach a confluency of around 50-70%. ER Tracker is added to the culture medium with a final concentration of 1 μM. Similarly, LDs are labeled by BODIPY and added to the culture medium with a final concentration of 5 μM. The cells are then incubated for 30 min at 37□°C and 5% CO_2_ concentration before RPOC. For BODIPY labeling, cells are washed twice with a warm culture medium after BODIPY incubation before RPOC.

## Statistics and reproducibility

The biological studies are repeated at least once for all experiments. Five cells are analyzed in each condition to obtain statistical results in **Fig. 2c, d**.

## Data availability statements

The authors confirm that the data supporting the findings of this study are available within the article [and/or] its supplementary materials. The data that support the findings of this study are available from the corresponding author, [author initials], upon reasonable request.

## Acknowledgment

This work is supported by NIH R35GM147092. The authors acknowledge Dr. Mingji Dai and Dr. Yiyang Luo for the synthesis of PST-1, which is published previously.

## Author contributions

BD and MGC built the confocal microscope and RPOC platform. KB built the galvo scanning system. BD and SHM performed measurements on biological samples and quantitative image analysis. SHM and SM cultured cells. SHM performed chemical treatment of cells. MC designed and built the comparator circuit. GC was involved in the design of the comparator circuit. KJM measured the RPOC system response time. CZ designed and supervised the project. CZ, BD, and SHM wrote the manuscript.

