## Supplementary Information for "Spatiotemporal precise optical manipulation of intracellular molecular activities"

**Supplementary Notes**

**1. Two-channel comparator circuitry**

The workflow of the two-channel comparator circuit is shown in **Fig. 1b**. A photograph of the circuit box is shown in **Supplementary Figure 1**. Two signal inputs can be connected to two separate detection channels and compared with preset thresholds. The threshold can be selected from a manual selection knob or input from separate BNC ports. Switches are used to select manual thresholds or digital input thresholds for signal comparison. If the intensity of the fluorescent signal exceeds the respective threshold, a '1' Transistor-Transistor Logic (TTL) command is sent to the output. Otherwise, a TTL ‘0’ is the output command. Invert functions can be selected to invert the output commands. This output can be directly sent to AOMs via comparator outputs to direct or stop the 1^st^ order laser beam to the sample. In this paper, thresholds are selected manually. Copies of the input thresholds are available as output.

In addition, logic gates are available to make logic calculations before sending the TTL to the AOM via logic outputs. AND, OR, and NOT functions are available separately for both of the two logic outputs. This function allows us to make higher-order decisions based on signals from two detectors. AOMs can also be controlled to be constantly on or constantly off by using AOM control switches before the logic outputs. Aside from output ports for AOM control, buffered output replicates the input signals which can be connected directly to the image acquisition channels to display images. Using a Tee connector at the TTL outputs, APXs can be directly visualized while driving AOM with TTL signals.

**Supplementary Figure 2a, b** shows two connection methods to achieve the same APX for both blue (405 nm) and green (532 nm) lasers for opto-control. **Supplementary Figure 2c** shows a fluorescence image of pollens excited using the 589 nm laser, and the corresponding APXs of the green and blue lasers selected using the connection in panel b. Note that both green and blue lasers can be simultaneously turned on only at the pollens. Here, only the input channel 1 is used. **Supplementary Figure 3a, b** shows the connection and selection of green APXs on pollens while blue APX outside pollens simultaneously. By flipping two switches at the inverters, the APXs from the green and blue lasers can be swapped (**Supplementary Figure 3c, d**). The thresholds for APXs from the two lasers are separately tunable if the input signal is split by a Tee connector and connected to both inputs.

**2. The response time of CW-RPOC**

The response time of the feedback system is measured using an oscilloscope. A signal pulse is sent from a function generator to both the oscilloscope and the comparator circuit. The latter generates a TTL ‘1’ signal that controls the AOM. The 1^st^ order laser from the AOM is acquired by a fast photodiode and sent to the oscilloscope. Comparing the time difference between the input signal and the response, the response time of the 405 nm laser is measured to be 580 ns (**Supplementary Figure 4a**). Similarly, the response time of the 532 nm laser is measured to be 658 ns (**Supplementary Figure 4b**). The delay is mostly generated by the AOM driver and AOM crystal response. Such a speed, however, is still much faster than the 10-20 µs pixel dwell time. The fast response time ensures simultaneous chemical detection and opto-control from the same pixel.

**3. The spatial resolution and opto-control precision of CW-RPOC**

The spatial resolution is characterized using a fluorescence image from HeLa cells transfected with EB3-EGFP (**Supplementary Figure 5a**). Fluorescence signals are excited using the 473 nm laser at 30 µW. A signal intensity line profile is plotted along a line that crosses one of the smallest structures in the image (**Supplementary Figure 5b**). A Gaussian fitting gives a full width at half maximum (FWHM) of ~309 nm. We, therefore, conclude that the spatial resolution of the CW-RPOC system using the 473 nm laser is 309 nm. All laser beams are collinearly combined and overfill the back aperture of the objective lens. Therefore, the spatial precision of RPOC using 532 and 405 nm wavelength lasers can be estimated to be 348 nm and 265 nm, respectively.

**4. Impact of 405 nm laser on tubulin dynamics studied by confocal fluorescence microscopy**

To further explore the impact of blue lasers on the cellular EB3-EGFP signals and the tubulin polymerization process, we used a commercial confocal fluorescence microscope (LSM510, Zeiss) for imaging and simultaneous laser treatment. Note that the commercial confocal microscope can perform simultaneous imaging and laser treatment but cannot selectively direct lasers solely at selected organelles as RPOC. Therefore, lasers are interacting with all pixels in the field of view (FOV). Cells are treated with 30 mW 405 nm laser for approximately 10 minutes. No significant fluorescence signal loss is detected (**Supplementary Figures 6a-c**) after treatment, but the tubulin polymerization process is stopped by the blue light. The 405 nm laser is not inducing detectable photobleaching of EB3-EGFP but largely impacting the microtubule dynamics.

**Supplementary Figures**

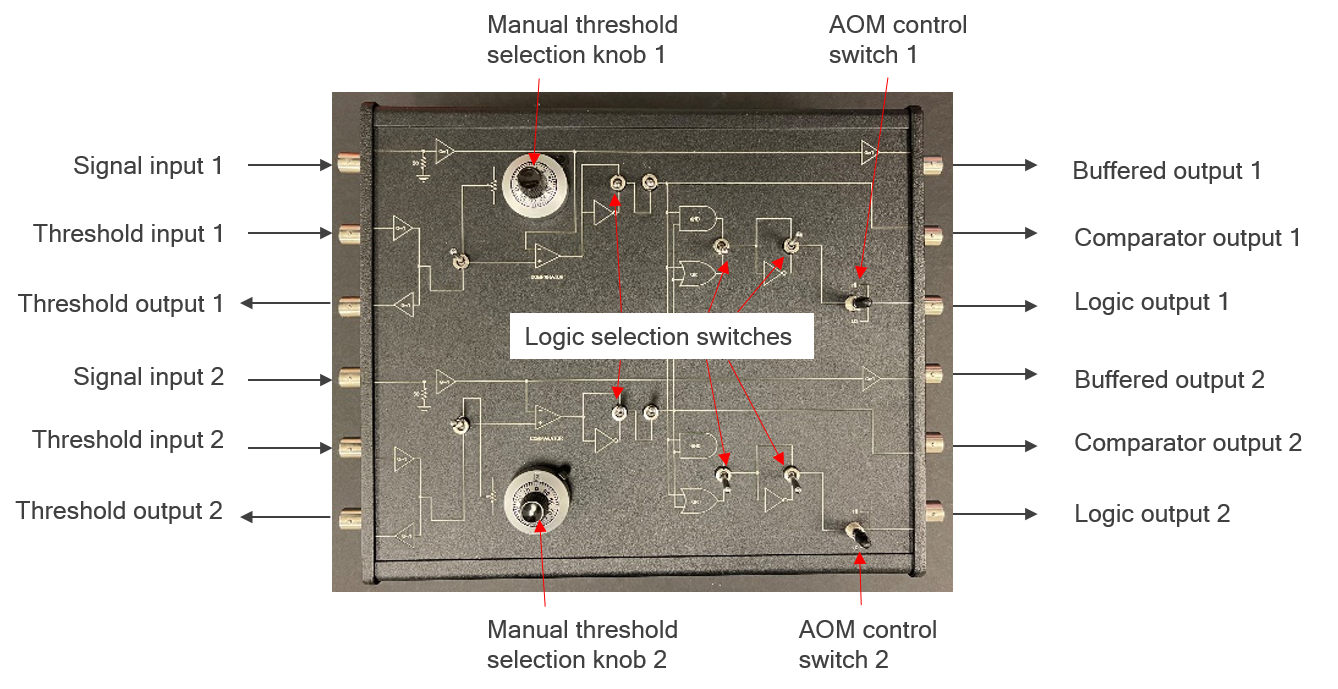

**Supplementary Figure 1.** A photograph shows the design of the two-channel comparator circuit box.

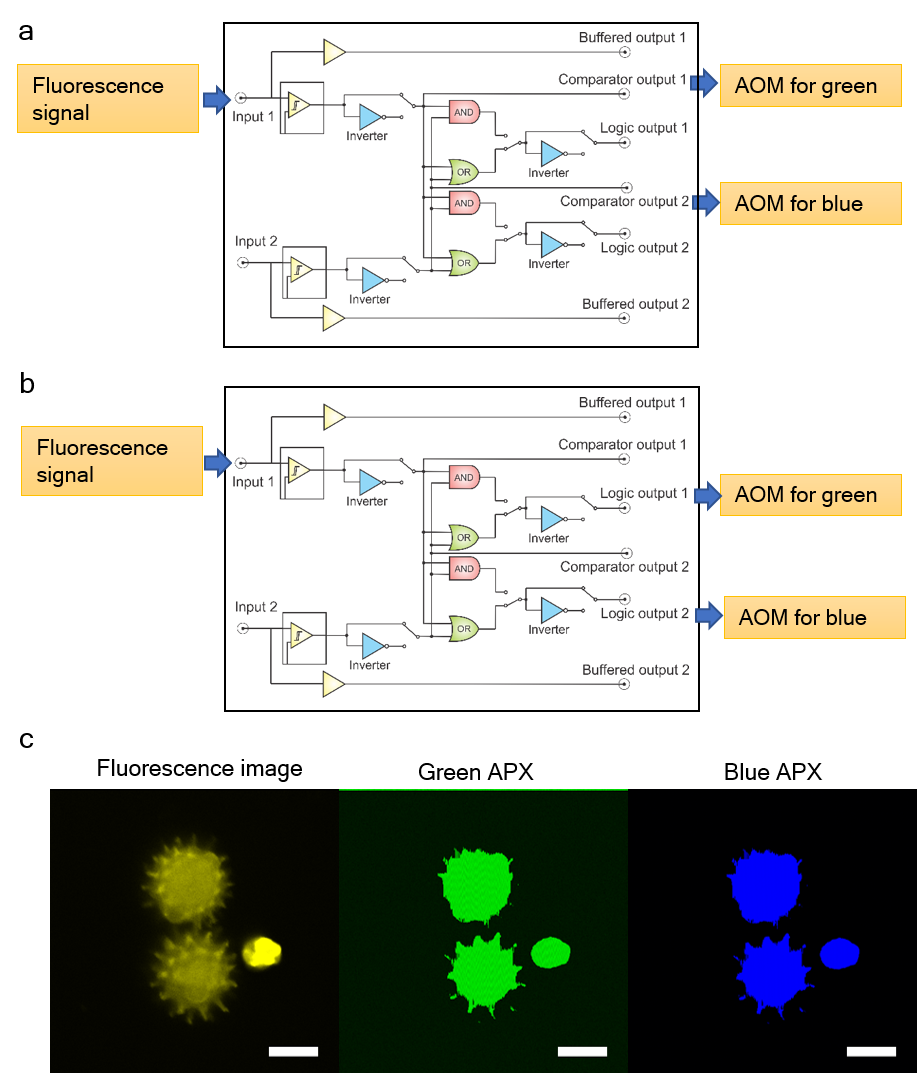

**Supplementary Figure 2.** (a, b) Two ways of controlling blue (405 nm) and green (532 nm) lasers separately using comparator circuit boxes. The APXs for blue and green lasers are selected for the same chemical contrast simultaneously. (c) A fluorescence image from pollens and the corresponding APXs for green and blue lasers selected from the same signal.

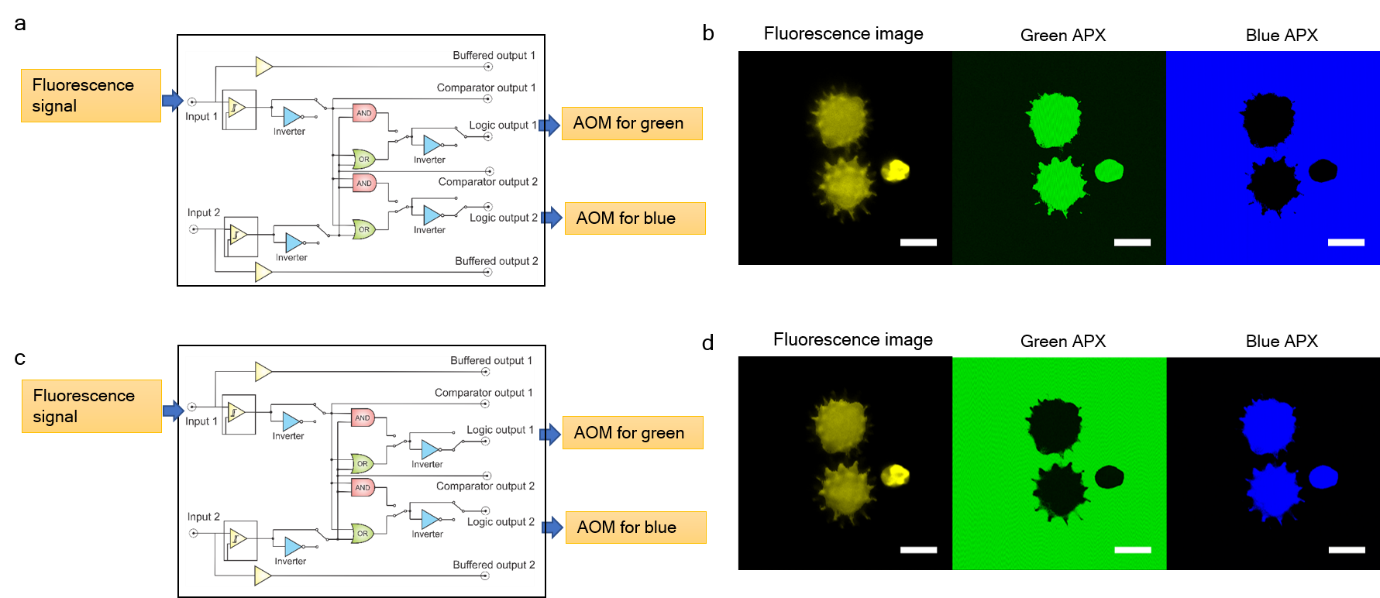

**Supplementary Figure 3.** (a) An example comparator circuit connection for achieving compensating APXs for blue and green lasers. (b) A fluorescence image from pollens and the corresponding APXs for green and blue lasers. The APXs for the green laser are selected on the pollen, while the APXs for the blue laser are selected outside the pollen simultaneously. (c, d) Similar to panels a and b, but invert APXs for blue and green.

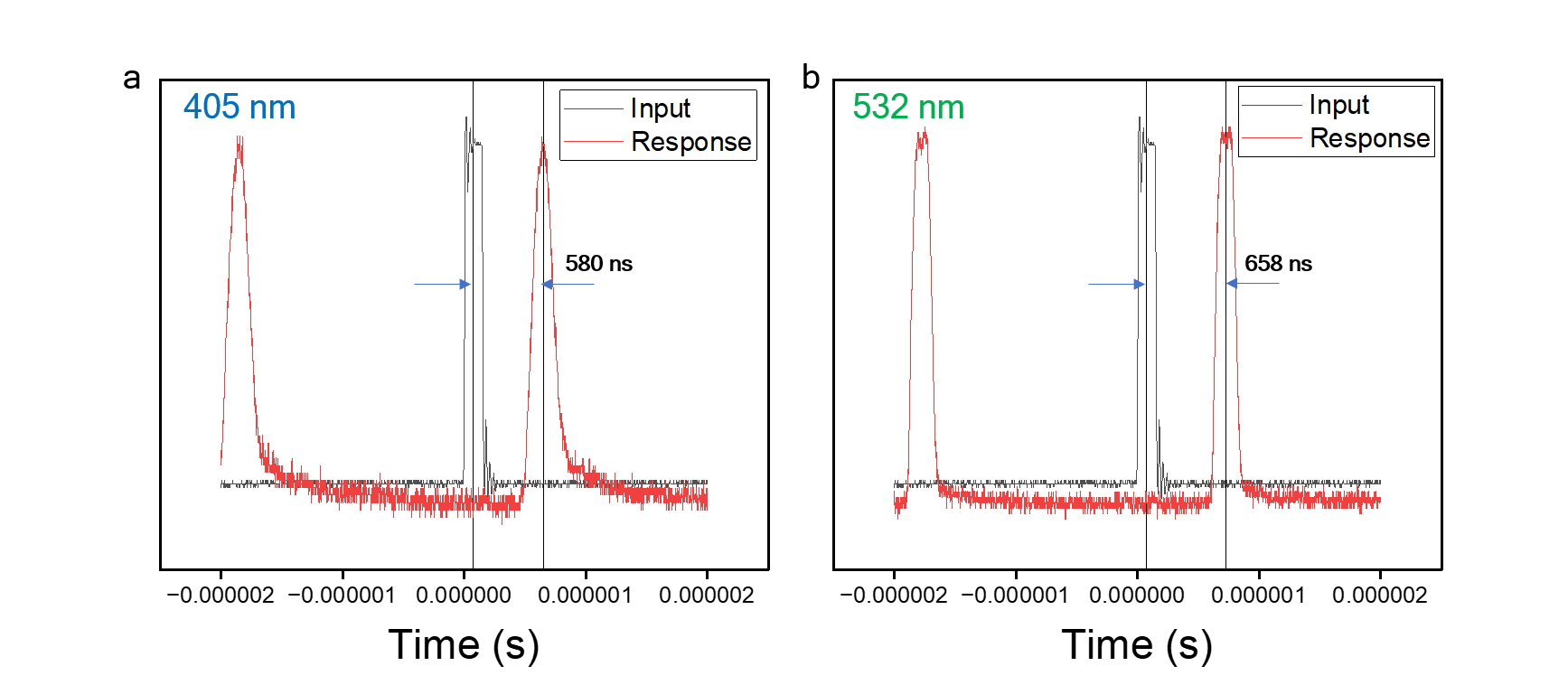

**Supplementary Figure 4.** (a) The response time of the RPOC feedback system is measured by a function generator and a photodiode for the 405 nm laser. (b) Similar to panel a, but for the 532 nm laser.

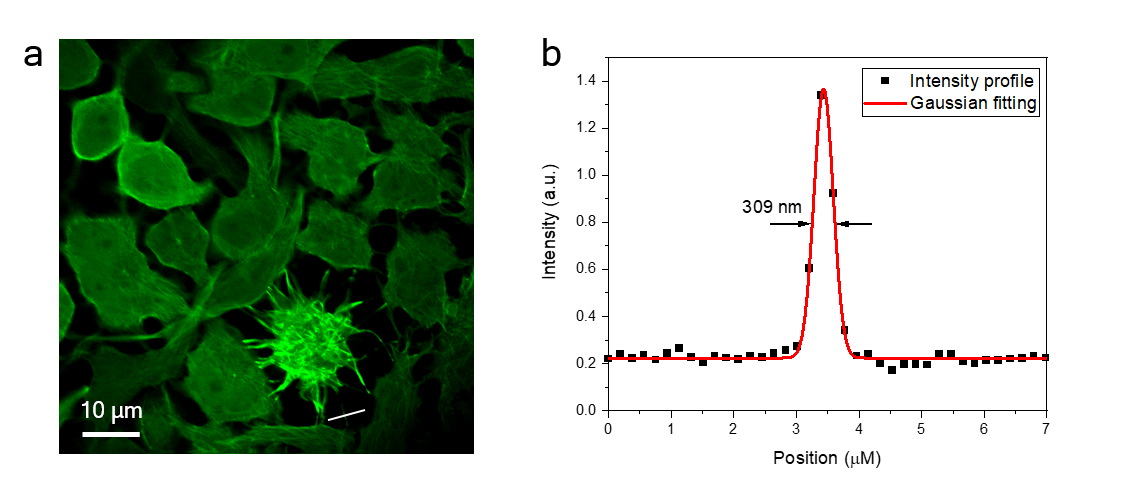

**Supplementary Figure 5.** (a) A fluorescence image of EB3-EGFP signals from HeLa cells. The images is averaged for 10 frames at 10 microsecond pixel dwell time. Signals are excited by the 473 nm laser. (b) The intensity profile along the line in panel a. Gaussian fitting gives a 309 nm spatial resolution.

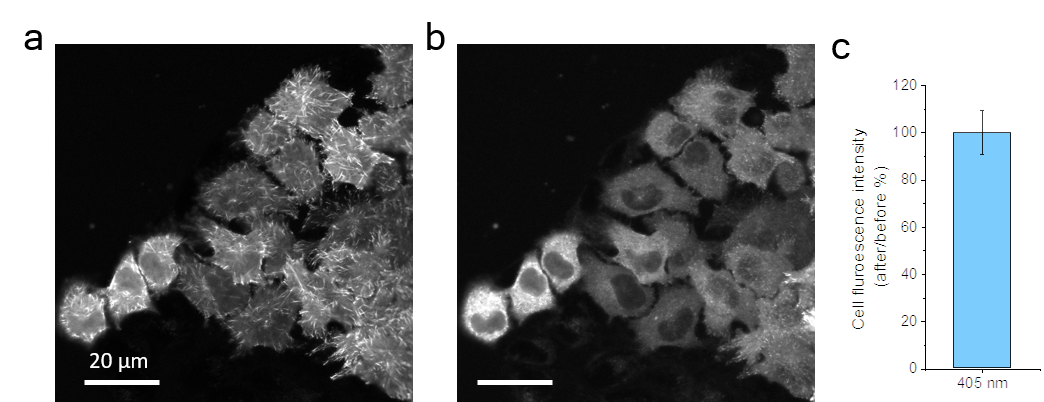

**Supplementary Figure 6.** (a) A fluorescence image of EB3-EGFP signals from HeLa cells acquired using a commercial confocal fluorescence microscope. (b) Image from the same FOV after 30 mW 405 nm Laser interaction throughout the FOV. (c) The fluorescence intensity ratio (I_after_/I_before_) from the same FOV after and before laser treatment.

**Supplementary Videos**

For all videos, the size of the field-of-views is 60 μm. The video consists of 30 frames with 400x400 pixels. The time interval between each frame is 5.2 s.

**Video captions**

**Video 1.1** Time-lapse EB3-EGFP signals in HeLa cells excited by 473 nm laser and detected by the CW-RPOC system.

**Video 1.2** Time-lapse APX intensity from 405 nm laser tracking high-intensity EB3 microtubule plus ends in HeLa cells.

**Video 1.2** Overlaid EGFP and APX time-lapse images of HeLa cells.

**Video 2.** Time-lapse EB3-EGFP signals from four laser interaction conditions in **Figure 2**.

**Video 3.** BODIPY signals from lipid droplets, APX selected from BODIPY signals, and the corresponding EB3-EGFP signals from HeLa cells during RPOC. The cells are treated with inactivated PST-1 before RPOC.

**Video 4.** ER-Tracker signals from lipid droplets, APX selected from ER-Tracker signals, and the corresponding EB3-EGFP signals from HeLa cells during RPOC. The cells are treated with inactivated PST-1 before RPOC.

**Videos 5, 6.** APXs from blue and green lasers at different parts of the HeLa cells and the corresponding EB3-EGFP signals from cells during RPOC. The cells are treated with inactivated PST-1 before RPOC.
